# Telomere-to-telomere gap-free genome assembly integrated with multi-omics uncovers shading mediated regulation of leaf aroma biosynthesis in aromatic crop *Pandanus amaryllifolius*

**DOI:** 10.64898/2026.03.12.711272

**Authors:** Zhijun Xu, Xuejiao Zhang, Yuzhan Li, Sheng Zhao, Qibiao Li, Jianan Zhou, Huan Ouyang, Xiaowen Hu

**Author notes:** These authors contributed equally to this study (Zhijun Xu, Xuejiao Zhang, Yuzhan Li, Sheng Zhao). Corresponding author at: Guangdong Modern Agriculture (Cultivated Land Conservation and Water-Saving Agriculture) Industrial Technology Research and Development Center, Zhanjiang Experiment Station, Chinese Academy of Tropical Agricultural Sciences, Zhanjiang 524013, China; South Subtropical Crops Research Institute, Chinese Academy of Tropical Agricultural Sciences, Zhanjiang 524091, China. E-mail address (X. Hu); (H. Ouyang); (J. Zhou); (Z. Xu).

## Abstract

*Pandanus amaryllifolius* is a valuable aromatic crop widely cultivated in tropical forest understories. Here, A telomere-to-telomere, gap-free genome assembly of the Chinese cultivar ‘Taishan Banlan’ was generated, and the molecular basis underlying shading-mediated regulation of leaf aroma biosynthesis was elucidated for the first time through integrated genomic, transcriptomic, and metabolomic analyses. Volatile metabolomics identified 1599 volatile organic compounds in leaf tissue, with terpenoids as the primary contributors to leaf aroma. Shading modulated the aroma quality of *P. amaryllifolius* primarily via terpenoid biosynthesis, yielding aroma profiles dominated by sweet, fruity, green, and woody notes. Integrated transcriptomic profiling revealed that terpenoid biosynthesis is co-regulated by structural genes in the MVA/MEP pathways, TPS family members, and key transcription factors, which collectively modulate the biosynthesis and accumulation of the three core differential terpenoids. Our findings provide valuable resources for the targeted genetic improvement and efficient cultivation optimization of *P. amaryllifolius*.

## 1. Introduction

*Pandanus amaryllifolius*, commonly known as Pandan or Banlan, is widely acclaimed as the "Oriental Vanilla" owing to its distinctive rice-like fragrance (Yan et al., 2025). As a tropical perennial plant of the *Pandanaceae* family, it is extensively cultivated in Southeast Asia and the tropical regions of southern China (Amnan et al., 2022). *P. amaryllifolius* is native to the Moluccas Islands of Indonesia (Wakte et al., 2009; Wakte et al., 2012), and was first introduced to mainland China during the 1950s–1960s (Hao, 2020), with its earliest distribution recorded in tropical and subtropical areas such as Hainan, Guangdong, Fujian, and Yunnan provinces. The fragrance of *P. amaryllifolius* is generally believed to originate primarily from its high concentration of 2-acetyl-1-pyrroline (2-AP), with levels over tenfold higher than those in fragrant rice (Routray et al., 2010; Wakte et al., 2012). For generations, *P. amaryllifolius* has been highly valued for its culinary applications, serving as a key spice and natural food colorant (Sidek et al., 2025; Wakte et al., 2012). Additionally, it is recognized as a traditional ethnic medicinal herb in folk medicine, attributed to its rich bioactive compounds with antiviral (Ooi et al., 2004) and antioxidant (Nor et al., 2008) properties, which are utilized for addressing numerous common ailments and exhibit potential for cancer therapy (Wang et al., 2024). For instance, gallic acid and cinnamic acid, isolated from its leaves, have been shown to inhibit the growth of breast cancer cells (Ghasemzadeh et al., 2013; Wang et al., 2024). In recent years, *P. amaryllifolius* has also been widely applied in the daily chemical industry for the manufacturing of high-value-added perfumes and other daily chemical products (Wang et al., 2024). Moreover, characterized by low pest and disease incidence, low management costs, and a prolonged harvesting period (exceeding ten years), *P. amaryllifolius* has also been widely cultivated as a promising intercropped aromatic crop in tropical regions of China and has emerged as a burgeoning industry boosting farmers’ income (Hao, 2020).

Despite increasing research interest in *P. amaryllifolius*, the core biological mechanisms governing its unique agronomic and metabolic traits remain incompletely elucidated. Notably, *P. amaryllifolius* is typically cultivated under diverse forest canopies, thriving in shaded environments and exhibiting tolerance to high humidity and elevated temperatures. It has a relatively low photosynthetic light saturation point (550–600 μmol/(m²·s)), while leaf aroma biosynthesis is tightly linked to shading (or light conditions) (Guo et al., 2020). Moderate shading is widely recognized as an effective strategy to enhance aromatic compound accumulation in diverse crops, including rice (Xie et al., 2021) and tea (Niu et al., 2025). Consistent with this general regulatory pattern, an artificial shading pot experiment conducted previously demonstrated that 30% and 60% relative shading significantly promote the accumulation of 2-AP and multiple volatile compounds in *P. amaryllifolius* leaves (Tang et al., 2020). In contrast, excessive shading exerts adverse impacts on the plant, most notably resulting in the deterioration of aroma quality. Plant aromas are predominantly composed of volatile organic compounds (VOCs), and over 1700 VOCs have been identified across more than 90 plant species to date (Chen et al., 2025). These metabolites are primarily categorized into three classes: volatile terpenoids (VTs), volatile phenylpropanoids/benzenoids (VPBs), and fatty acid-derived volatiles (FADVs) (Fu et al., 2017; Zhou et al., 2023a). For *P. amaryllifolius*, its characteristic rice-like fragrance is directly attributed to 2-AP, and its VOC profile is further composed of furans, pyrroles, terpenes, alcohols, alkenes, and esters (Wakte et al., 2012; Yan et al., 2025). However, under shading conditions, whether the altered leaf aroma stems from fluctuations in 2-AP levels or variations in other VOCs remains unclear. The underlying molecular regulatory networks and metabolic pathways linking light intensity to aroma biosynthesis have also not been systematically characterized.

High-quality contiguous genome assemblies provide a foundational framework for investigating species evolution and deciphering the genetic and molecular underpinnings of desirable traits (Nunn et al., 2022; Yadav et al., 2025). The recently released draft genome of the Malaysian *P. amaryllifolius* cultivar ‘Sungai Buloh’ spans ∼494.1 Mb with 1448 contigs, which does not meet reference-quality standards (Sidek et al., 2025). Furthermore, functional annotation of its genes, particularly those related to 2-AP biosynthesis and other secondary metabolic pathways, remains insufficient and demands further investigation. Recent advancements in sequencing technologies, including long-read sequencing and high-throughput chromosome conformation capture (Hi-C) technology, alongside optimized assembly tools, have enabled the generation of telomere-to-telomere (T2T) genome assemblies for numerous species. Notable examples include *Arabidopsis thaliana* (Wang et al., 2022), *Oryza sativa* (Song et al., 2021), *Zea mays* (Chen et al., 2023a), *Glycine max* (Jia et al., 2024), *Linum usitatissimum* (Yadav et al., 2025), and many other crop species. Unlike conventional genome assemblies, T2T gap-free assemblies allow for the complete capture of key complex repetitive regions such as centromeres and telomeres, thereby facilitating the systematic analysis of non-coding and repetitive DNA elements. Such capabilities are indispensable for deciphering genome evolution and the key molecular mechanisms governing phenotypic variation (Yadav et al., 2025).

In view of the aforementioned knowledge gaps, the present study aimed to assemble a high-quality genome of *P. amaryllifolius* and utilize it to dissect the molecular mechanisms underlying shading-mediated regulation of aroma biosynthesis. To this end, we generated the first T2T gap-free genome of the Chinese Pandan cultivar ‘Taishan Banlan’ by integrating Oxford Nanopore Technology (ONT), PacBio HiFi, Hi-C, and next-generation sequencing (NGS) technologies. We performed comprehensive gene annotation and comparative genomic analyses to investigate its evolutionary history and characterize gene family expansions/contractions. Additionally, we conducted integrative volatile metabolomics and transcriptomics analyses to decipher how light intensity (shading) regulates volatile accumulation and leaf aroma biosynthesis in this species, with a specific focus on regulatory networks underlying terpenoid biosynthesis. Collectively, this study aimed to address the existing knowledge gaps, deliver a high-quality comprehensive genomic resource for *P. amaryllifolius*, and lay a solid theoretical foundation for its targeted genetic improvement and efficient cultivation optimization.

## 2. Materials and Methods

### 2.1 Plant material and genome sequencing

Fresh young leaves of *P. amaryllifolius* landrace ‘Taishan Banlan’ were collected for genome sequencing. This landrace was cultivated in the experimental field of Zhanjiang Experimental Station, Chinese Academy of Tropical Agricultural Sciences (20°9′50″ N, 110°15′42″ E). Flow cytometry, molecular karyotyping, and fluorescence in situ hybridization (FISH) analyses were performed following previously established protocols (Cápal et al., 2023). High-quality genomic DNA was isolated from *P. amaryllifolius* using the CTAB method. The genome was sequenced by BerryGenomics Co., Ltd. (Beijing, China) with an integrated sequencing strategy incorporating ONT ultra-long reads, PacBio HiFi reads, Hi-C reads, and NGS short reads. Specifically, ONT libraries were constructed using the Ligation Sequencing Kit 1D (SQK-LSK109, Oxford Nanopore Technologies, UK) and sequenced on the PromethION 48 platform. PacBio HiFi libraries (∼15 Kb) were prepared with the SMRTbell® Prep Kit 3.0 (102-182-700, PacBio, USA) and sequenced on the PacBio Revio platform. Hi-C and NGS libraries were generated following standard protocols and sequenced on the Illumina NovaSeq 6000 platform with 150 bp paired-end reads.

For RNA sequencing, five tissues (leaves, stems, roots, aerial roots, and lateral shootlets) subjected to different treatments (abscisic acid (ABA), ethylene (ET), jasmonic acid (JA), salicylic acid (SA), brassinosteroid (BR), polyethylene glycol (PEG), sodium chloride (NaCl), and sodium bicarbonate (NaHCO₃)) were collected. Total RNA was extracted using the Polysaccharide-Polyphenol Plant Total RNA Extraction Kit (TIANGEN, Beijing, China). For treated samples, RNA was isolated at 6 hours post-treatment. RNA-seq libraries were constructed following standard protocols and sequenced on the Illumina NovaSeq 6000 platform, generating 150-bp paired-end reads. For full-length transcriptome sequencing, total RNA isolated from the five aforementioned tissues was pooled at an equimolar ratio for library preparation. Sequencing libraries were constructed using the SMRTbell® Prep Kit 3.0 (102-182-700, PacBio, USA) and subsequently sequenced on the PacBio Revio platform.

### 2.2 Genome assembly and quality assessment

Initial estimation of the *P. amaryllifolius* genome characteristics was conducted via 21-k-mer frequency analysis using Jellyfish (v2.3.0) (Marçais et al., 2011). Subsequently, the k-mer distribution was analyzed with GenomeScope2 (v2.0) to determine genome size, heterozygosity, and repeat content (Ranallo-Benavidez et al., 2020). The genome assembly was generated using Hifiasm (v0.19.9) with the integration of PacBio HiFi and Hi-C sequencing data (Cheng et al., 2021). Hi-C reads were processed using HiC-Pro (v3.1.0) to retrieve valid read pairs (Servant et al., 2015); subsequently, contigs were anchored and ordered into chromosome-scale scaffolds via YaHS (v1.2) (Zhou et al., 2023b), with Hi-C contact frequency matrices serving as the reference. ONT reads were used to fill residual gaps in the assembly through TGS-GapCloser (v1.2.1) (Xu et al., 2020). Finally, the Hi-C contact heatmap was constructed using HapHiC (v1.0.6) (Zeng et al., 2024).

Assembly completeness was assessed using BUSCO (v5.7.1) with the embryophyta_odb10 database (Simão et al., 2015). Consensus quality value (QV) and k-mer completeness were evaluated via Merqury, which compares the assembled genome to the k-mer dataset derived from Illumina short reads (Rhie et al., 2020). For long terminal repeat (LTR) retrotransposon-related assembly quality, the LTR Assembly Index (LAI) was computed to serve as an assessment metric, utilizing the LTR_retriever (v2.9.0) software (Ou et al., 2018). Genome comparison between Chinese ‘Taishan Banlan’ and Malaysian ‘Sungai Buloh’ was performed using the GenomeSyn software (Zhou et al., 2022).

### 2.3 Genome annotation and repeat sequence identification

Repetitive sequences in the *P. amaryllifolius* T2T genome were predicted using a combined strategy of *de novo* annotation and homology-based searching. Transposable elements (TEs) were identified by MITE-Hunter (Han et al., 2010), EDTA (v2.2.1) (Su et al., 2021), RepeatModeler (v2.0.4) (Flynn et al., 2020), LTR_FINDER (Ou et al., 2019), LTRharvest (Ellinghaus et al., 2008), and LTR_retriever (v2.9.0) (Ou et al., 2018). Tandem repeats were predicted using TRF (v4.09) (Benson, 1999) and Krait (v1.5.1) (Du et al., 2018). RepeatMasker was employed to screen the genome sequences homologous to those in the RepBase database (http://www.girinst.org/repbase), followed by masking of repetitive regions in the genome based on this integrated repeat library. Subsequently, *de novo* annotation was performed on the repeat-masked genome using RepeatModeler to identify additional uncharacterized repetitive sequences.

Gene structure annotation of the *P. amaryllifolius* T2T genome was conducted via a multi-pronged strategy, including *de novo* prediction, transcriptome-supported prediction, and homology-based prediction (with reference to *Ananas bracteata*, *Ananas comosus*, *Asparagus setaceus*, *Dioscorea alata*, *Dioscorea cambodiana*, *Oryza sativa*, and *Arabidopsis thaliana*). Prediction results from these approaches were integrated using EvidenceModeler to generate a consensus gene structure annotation (Haas et al., 2008). Gene functions were annotated based on sequence similarity to entries in multiple public databases: Gene Ontology (GO), Kyoto Encyclopedia of Genes and Genomes (KEGG), NCBI non-redundant protein database (Nr), Pathway, Universal Protein Resource (UniProt), EuKaryotic Orthologous Groups (KOG), and InterPro.

Non-coding RNA (ncRNA) annotation was performed using tRNAscan-SE (Chan et al., 2019) and the Rfam database (Ontiveros-Palacios et al., 2025). Specifically, tRNAscan-SE was used to predict tRNAs in the genome, while other ncRNA types (rRNAs, miRNAs, snRNAs, and snoRNAs) were annotated by BLAST against the Rfam database.

### 2.4 Comparative and evolutional analysis of *P. amaryllifolius*

Eleven publicly available genome assemblies (i.e., *Ananas bracteatus*, *Amborella trichopoda*, *Ananas comosus*, *Arabidopsis thaliana*, *Asparagus setaceus*, *Dioscorea alata*, *D. cambodiana*, *Durio zibethinus*, *Oryza sativa*, *Solanum lycopersicum*, and *Zostera marina*), were downloaded to conduct comparative and evolutionary analyses of *P. amaryllifolius*. Protein sequences were extracted from *P. amaryllifolius* and the 11 aforementioned genomes, and subjected to OrthoFinder (v2.4) (Emms et al., 2019) for orthologous groups identification. Single-copy orthologous genes were retrieved from all these 12 species and subjected to reciprocal all-versus-all alignment using MAFFT (v7.205) (Katoh et al., 2013). A phylogenetic tree of these 12 species was constructed with IQ-TREE (v1.6.11) (Nguyen et al., 2015) and calibrated using fossil records retrieved from the TimeTree database (https://timetree.org/). CAFE (v5.1) software was utilized to infer gene family contraction and expansion based on the orthologous group annotations and the constructed phylogenetic tree (Mendes et al., 2021). Functional enrichment analysis of contracted and expanded gene families was performed using the KEGG and GO databases. Whole-genome duplication (WGD) events were detected via Ks analysis of collinear gene pairs using WGDI tool (v0.75) (Sun et al., 2022), and density plots for different species were generated with ggplot2 (v2.2.1) (https://ggplot2.tidyverse.org/). LTR insertion times were estimated using EDTA package (v2.2.2), following the method described by Yang et al. (2024). Additionally, collinearity analysis between *P. amaryllifolius* and two reference species (*D. alata* and *A. thaliana*) was conducted using TBtools software (v2.390) (Chen et al., 2023b). The karyotype evolution was analyzed based on the collinearity relationship of ancestral eudicot karyotypes (AEK) with *P. amaryllifolius* using WGDI (Sun et al., 2022).

### 2.5 Leaf sample preparation under saturating light and 50% of saturating light

One-year-old tissue-cultured *P. amaryllifolius* seedlings were selected as experimental materials. The seedlings were transplanted into plastic pots (12 cm (diameter) × 15 cm (height)) filled with a growth substrate composed of nutrient soil and vermiculite mixed at a 3:1 (v/v) volume ratio, and acclimatized for one month. Exterior leaves of the seedlings were then removed, with five young heart leaves retained per seedling. Two light intensity treatments were eatablished in a plant growth chamber: saturating light (100%) and 50% of saturating light. Using the pre-determined seedling light saturation point (600 μmol photons·m⁻²·s⁻¹) (Guo et al., 2020) and conversion factor (1 klux = 12.5 μmol photons·m⁻²·s⁻¹), target illuminances were calculated as 48 klux (saturating light) and 24 klux (50% saturating light). Additional growth chamber environmental parameters were standardized: temperature (25°C), relative humidity (80%), a 16-h light/8-h dark photoperiod, and uniform soil moisture. Sampling was conducted at 1, 3, and 7 weeks after the initiation of light treatment. Two outermost leaves were harvested per seedling,and leaves from three seedlings were pooled as one biological replicate. Each replicate was divided into three aliquots: one for total RNA extraction, one for volatile aroma metabolite profiling, and one for 2-AP content quantification in leaves. Three independent biological replicates were set up per treatment, and all aliquots were immediately flash-frozen in liquid nitrogen and stored at −80°C.

### 2.6 GC-MS analysis of VOCs in leaves under saturating light and 50% of saturating light

To characterize the VOCs profile of *P. amaryllifolius*, solid phase microextraction-gas chromatography-mass spectrometry (SPME-GC-MS) analyses were conducted by Metware Biotechnology Co., Ltd. (Wuhan, China). Leaf samples were ground to a fine homogeneous powder under liquid nitrogen, and 0.5 g aliquots of the powder were transferred to 20 mL headspace vials (Agilent, USA) for VOC analysis. SPME extraction conditions, as well as VOC identification and quantification, were carried out following the method previously described by Chen et al. (2025). The determination of total 2-AP content in leaves was carried out according to the protocol reported by Zhang et al. (2023).

Principal component analysis (PCA), orthogonal partial least squares-discriminant analysis (OPLS-DA), and hierarchical clustering analysis (HCA) were conducted using the MetaboAnalyst package (Chong et al., 2018; Chong & Xia, 2018). Variable importance in projection (VIP) values (VIP ≥ 1) and metabolite fold change (FC) (|Log₂FC| ≥ 1) were used to identify differentially accumulated volatiles (DAVs). These DAVs were annotated using the KEGG COMPOUND Database (http://www.kegg.jp/kegg/compound/) and further mapped to KEGG PATHWAY (http://www.kegg.jp/kegg/pathway.html) for pathway enrichment analysis. Additionally, sensory odor characterization of the identified leaf VOCs was performed using public online databases: http://www.thegoodscentscompany.com, http://www.odour.org.uk/odour/index.html, http://perflavory.com/, and http://foodflavorlab.cn/.

### 2.7 Transcriptome sequencing of leaf samples under saturating light and 50% of saturating light

Total RNA extraction, quality control, library construction, and sequencing were performed as described in Section 2.1. Raw read quality control and filtering, clean read alignment to the genome, and novel transcript assembly were performed using FastQC (https://www.bioinformatics.babraham.ac.uk/projects/fastqc/), fastp (v0.39) (Chen et al., 2018), HISAT2 (v2.1.0) (Kim et al., 2019), and StringTie (v1.3.6) (Pertea et al., 2015), respectively. Gene expression levels were measured and normalized as fragments per kilobase of transcript per million fragments mapped (FPKM) values. Differentially expressed genes (DEGs) were identified using DESeq2 (Love et al., 2014) with thresholds of |log₂FoldChange| ≥ 1 and false discovery rate (FDR) ≤ 0.05. The functions of DEGs were annotated via KEGG and GO enrichment analyses. Weighted gene co-expression network analysis (WGCNA) was conducted via Metware Cloud (https://cloud.metware.cn) using the FPKM values of the identified DEGs. The power parameter was estimated with an R² cutoff value of 0.85. Co-expression modules were defined using the dynamic tree cut algorithm, with a minimum module size of 50 and a merge cut height of 0.25 for similar modules.

### 2.8 Analysis of genes related to terpenoid biosynthesis and terpene synthases

Terpenoid biosynthesis-related genes in the mevalonate (MVA) and methylerythritol phosphate (MEP) pathways were identified from the *P. amaryllifolius* genome using genome annotation and sequence similarity to *A. thaliana* proteins. Terpene synthase (TPS) members in *P. amaryllifolius* were identified via two approaches: BLASTP searches against *A. thaliana* TPS proteins and hidden Markov model (HMM) searches with conserved Pfam domains (PF01397 and PF03036). A phylogenetic tree of TPS members from different species was constructed using MEGA (v7.0) (Kumar et al., 2016). TPS members were classified based on their phylogenetic relationships with *A. thaliana* TPS proteins and previous classification systems (Aubourg et al., 2002; Song et al., 2020). Gene expression heatmap and collinearity analyses of TPS family members among *P. amaryllifolius*, *A. thaliana*, and *D. alata* were performed using TBtools (Chen et al., 2023b). A co-expression network involving terpenoid biosynthesis structural genes, TPS genes, and transcription factors (TFs) was constructed with Cytoscape (v3.8.5) (Shannon et al., 2003). The relative expression levels of these genes were validated via qRT-PCR, with all primers used in this study listed in Table S1.

## 3 Results

### 3.1 Genome assembly and quality assessment

The *P. amaryllifolius* landrace ‘Taishan Banlan’ was selected as the reference material for genome sequencing and assembly (Fig. 1a; Fig. S1). Cytogenetic analyses were performed on *P. amaryllifolius* accessions collected from three geographically distinct provinces in China (Guangdong, Yunnan, and Hainan), and the results showed that all tested samples were diploid (2n = 60). Four distinct 45S rDNA hybridization signals were detected in these samples using a 45S rDNA probe (Fig. 1b, c; Fig. S2). Genome size was estimated by flow cytometry and k-mer-based analysis, yielding estimates ranging from 468.35 to 481.76 Mb (Fig. S3).

**Fig. 1.**
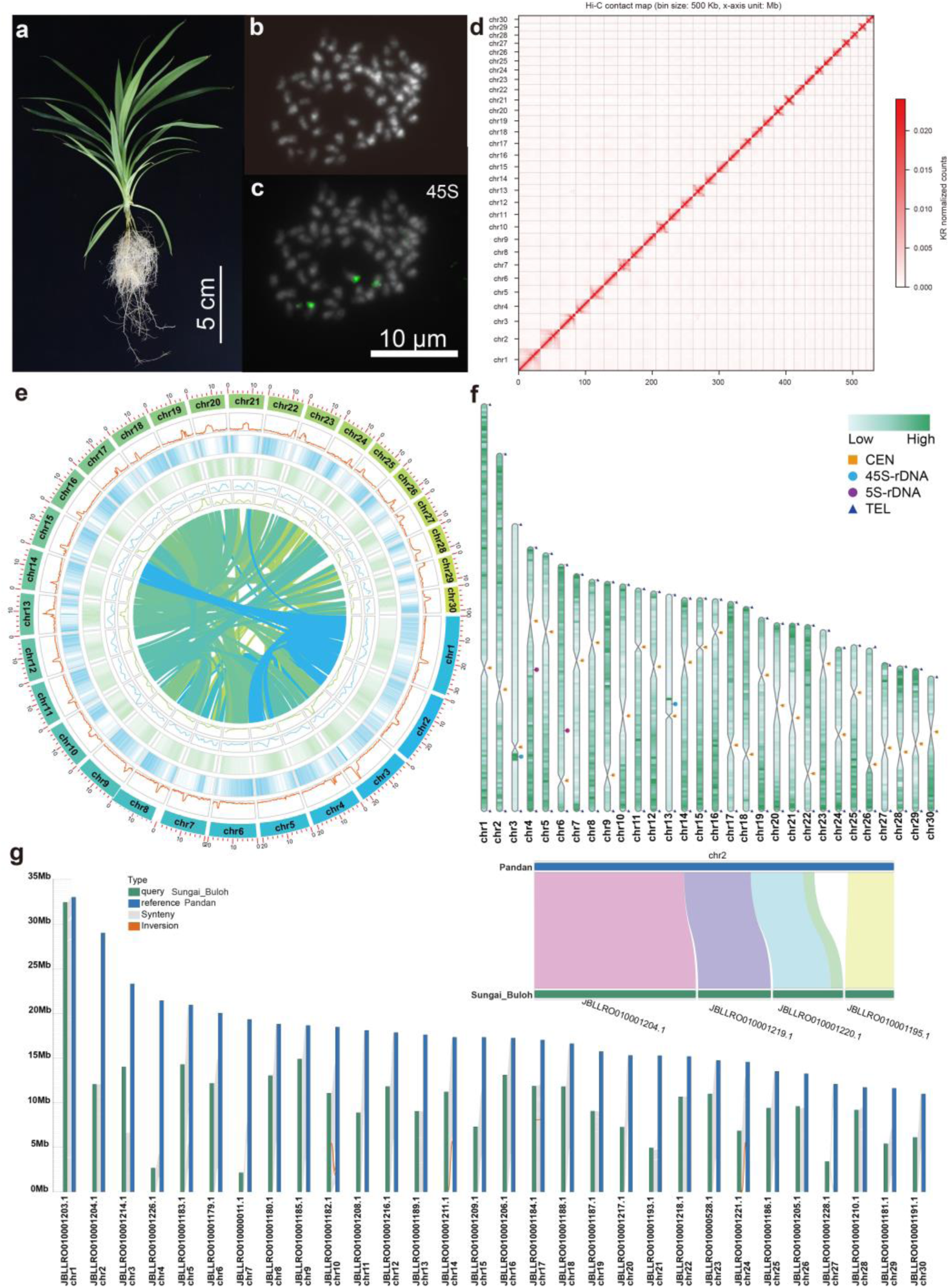
Genome assembly of *P. amaryllifolius*. (a) *P. amaryllifolius* landrace ‘Taishan Banlan’; (b–c) Karyotype and FISH analyses of *P. amaryllifolius*; (d) Hi-C interaction map of *P. amaryllifolius*; (e) Circos plot showing the genomic landscape of *P. amaryllifolius* genome (circles from innermost to outermost represent the gene collinearity, Gypsy density, Copia density, GC content, repeat density and gene density); (f) Distribution of telomere, centromere, 5S-rDNA and 45S-rDNA on *P. amaryllifolius* chromosomes; (g) Genome synteny between the present *P. amaryllifolius* assembly and the Malaysian cultivar ‘Sungai Buloh’.

To generate a high-quality reference genome, the *P. amaryllifolius* genome was sequenced using ONT ultra-long, PacBio HiFi, Hi-C, and NGS platforms, resulting in 32.85 Gb (∼62.52×), 38.02 Gb (∼72.35×), 52.91 Gb (∼100.69×), and 84.76 Gb (∼161×) of clean reads, respectively (Table S2–Table S5). The genome was assembled using the ONT ultra-long and PacBio HiFi data, and successfully anchored to 30 pseudochromosomes based on the Hi-C assembly, matching the cytogenetically confirmed chromosome count (Fig. 1d). After successive rounds of polishing, gap filling, and error correction, a final T2T, non-redundant, and gap-free genome assembly was generated, with a total length of 525.46 Mb (Table 1, Table S6).

**Table 1.**
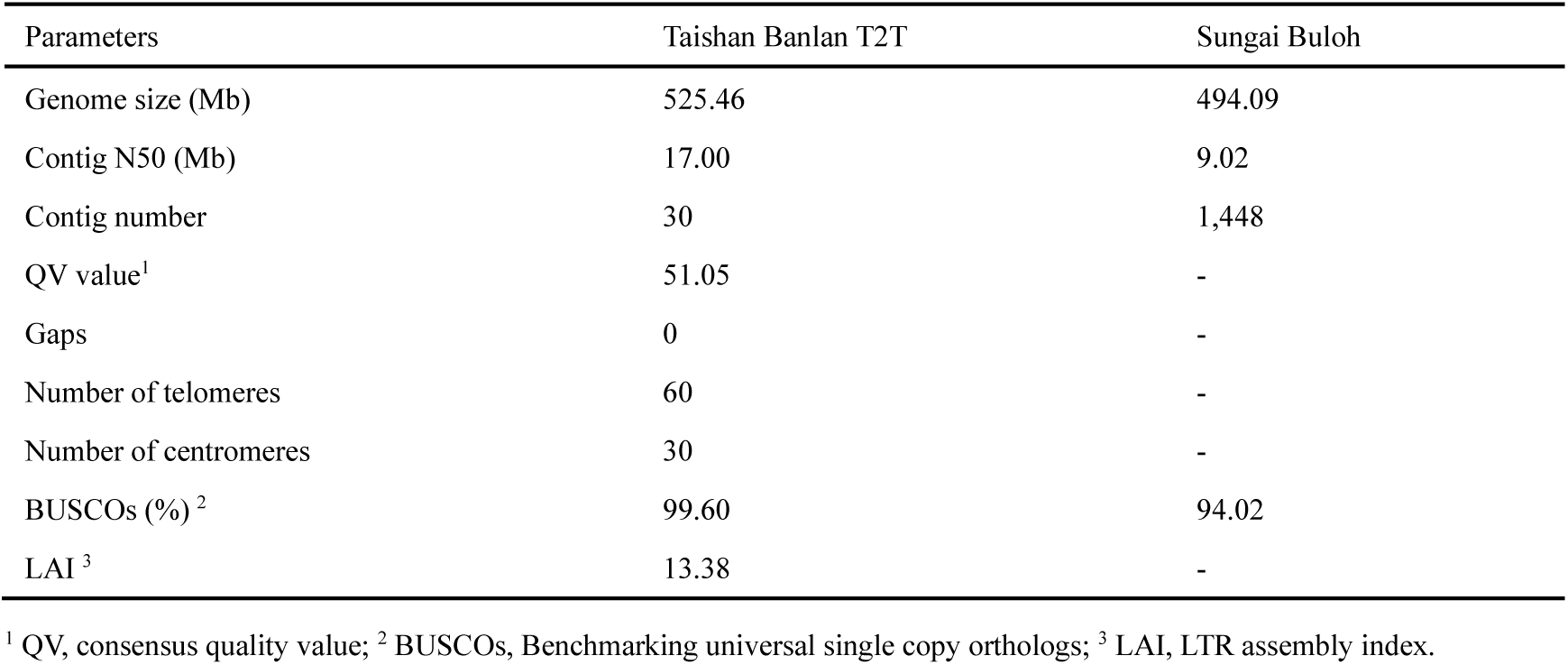
Summary of T2T gap-free genome assembly of *P. amaryllifolius*.

| Parameters | Taishan Banlan T2T | Sungai Buloh |
| --- | --- | --- |
| Genome size (Mb) | 525.46 | 494.09 |
| Contig N50 (Mb) | 17.00 | 9.02 |
| Contig number | 30 | 1,448 |
| QV value<sup>1</sup> | 51.05 | - |
| Gaps | 0 | - |
| Number of telomeres | 60 | - |
| Number of centromeres | 30 | - |
| BUSCOs (%)<sup>2</sup> | 99.60 | 94.02 |
| LAI<sup>3</sup> | 13.38 | - |

<sup>1</sup> QV, consensus quality value; <sup>2</sup> BUSCOs, Benchmarking universal single copy orthologs; <sup>3</sup> LAI, LTR assembly index.

To evaluate the accuracy and consistency of the T2T assembly, sequencing reads from all platforms were mapped back to the T2T genome. High alignment rates were achieved: 98.35% of Illumina short reads, 99.99% of PacBio HiFi reads, and 99.96% of ONT reads mapped successfully (Table S7). The QV of the genome was estimated to be 51.05, indicating a low base-level error rate across chromosomes (Table S8). BUSCO assessment revealed that the T2T genome exhibited a high completeness of 99.6%, demonstrating excellent assembly quality (Table S9). The LAI of the genome was 13.38, meeting the reference genome standard. Compared with the genome assembly of the Malaysian cultivar ‘Sungai Buloh’, the present *P. amaryllifolius* assembly exhibits superior chromosomal continuity and completeness (Fig. 1g and Fig. S4). Collectively, these metrics demonstrate that the *P. amaryllifolius* T2T genome assembly achieves high completeness, accuracy, and structural integrity.

### 3.2 Characterization and annotation of centromeres and telomeres

With the gap-free genome assembly available, centromeric and telomeric regions were subsequently characterized to elucidate chromosome-scale structural organization. Tandem repeat annotation coupled with structural hierarchy analysis identified 64-bp and 144-bp sequences as the predominant repeat monomers genome-wide (Fig. S5). However, no single monomer sequence showed strict chromosome-specific enrichment, suggesting a heterogeneous centromeric repeat composition in *P. amaryllifolius*. Putative centromeric regions were systematically inferred across all 30 chromosomes by integrating multiple genomic features, including 21-k-mer density, gene length distribution, LTR length distribution, tandem repeat abundance, and Hi-C intrachromosomal interaction patterns (Fig. 1f, g; Fig. S6; Table S10). These candidate regions exhibited substantial size variation, ranging from 0.48 to 3.96 Mbp. Notably, the inferred centromeres of chr22 and chr23 localized near the chromosomal termini, indicating non-canonical centromere positions. The canonical telomere repeat monomer was identified as AAACCCT/AGGGTTT, and 60 telomeric sites were detected across the entire genome (Fig. 1f). The copy number of telomeric repeat monomers ranged from 63 to 1831 among the 30 chromosomes, reflecting substantial variation in telomere length (Table S11).

### 3.3 Genome annotation

Following integration and removal of redundant annotations, a total of 304.10 Mb of repetitive sequences were identified, accounting for 57.87% of the assembled genome. Among these, LTR retrotransposons were the dominant repeat class, representing 34.52% of the genome. The two major LTR subfamilies were LTR-Copia (8.79%) and LTR-Gypsy (8.08%), respectively (Fig. 1e; Table S12). In contrast, DNA transposons were comparatively scarce, accounting for only 1.98% of the genome. Tandem repeats contributed an additional 6.21% of the genome, including microsatellites (0.48%), minisatellites (2.08%), and satellite repeats (3.67%) (Table S13). Overall, these findings demonstrate that the *P. amaryllifolius* genome is highly enriched in retrotransposon-derived sequences, consistent with its large genome size and high repeat content.

Gene structure prediction was performed on the repeat-masked genome assembly, yielding 23,819 protein-coding genes, with an average gene length of 8,499.72 bp and 6.51 exons per gene (Table S14). Of these, 22,066 genes (92.64% of the predicted protein-coding genes) were annotated in at least two databases (Fig. S7; Table S15), indicating broad coverage of conserved plant gene functions. In addition, ncRNAs were systematically annotated, yielding 2,011 rRNAs, 148 miRNAs, 359 tRNAs, 94 snRNAs, and 724 snoRNAs (Table S16). Further analysis of 5S rDNA and 45S rDNA revealed large rRNA clusters: 5S rDNA clusters were localized on chr4 (9.85–10.05 Mb) and chr6 (13.45–13.49 Mb), whereas 45S rDNA clusters were detected on chr3 (18.52–19.19 Mb) and chr13 (9.39–9.40 Mb). These results offer comprehensive insights into the repeat composition and gene architecture of the *P. amaryllifolius* genome, laying a solid foundation for subsequent functional and comparative genomic studies.

### 3.4 Comparative genomics and evolutionary history of *P. amaryllifolius*

To examine the genomic characteristics and evolutionary position of *P. amaryllifolius*, comparative genomic analyses were performed with 11 other representative plant species covering both monocots and dicots. A total of 380,350 orthologous gene families were identified, of which 21,859 genes (91.91%) of *P. amaryllifolius* were clustered into 12,150 orthologous groups (3.19%) (Table S17), indicating broad conservation of gene content. A core set of 7,733 orthologous gene families was universally conserved across all 12 species (Fig. 2a). Notably, *P. amaryllifolius* possessed 246 species-specific orthologous groups comprising 363 paralogous genes, corresponding to the fewest species-specific orthologous groups among all tested species (Fig. 2a, 2b). Functional enrichment analysis revealed that these species-specific genes were predominantly enriched in biological processes involved in transcription and RNA metabolism regulation, biosynthesis and metabolism regulation, stress response, and global biological process regulation (Fig. S8, Fig. S9).

**Fig. 2.**
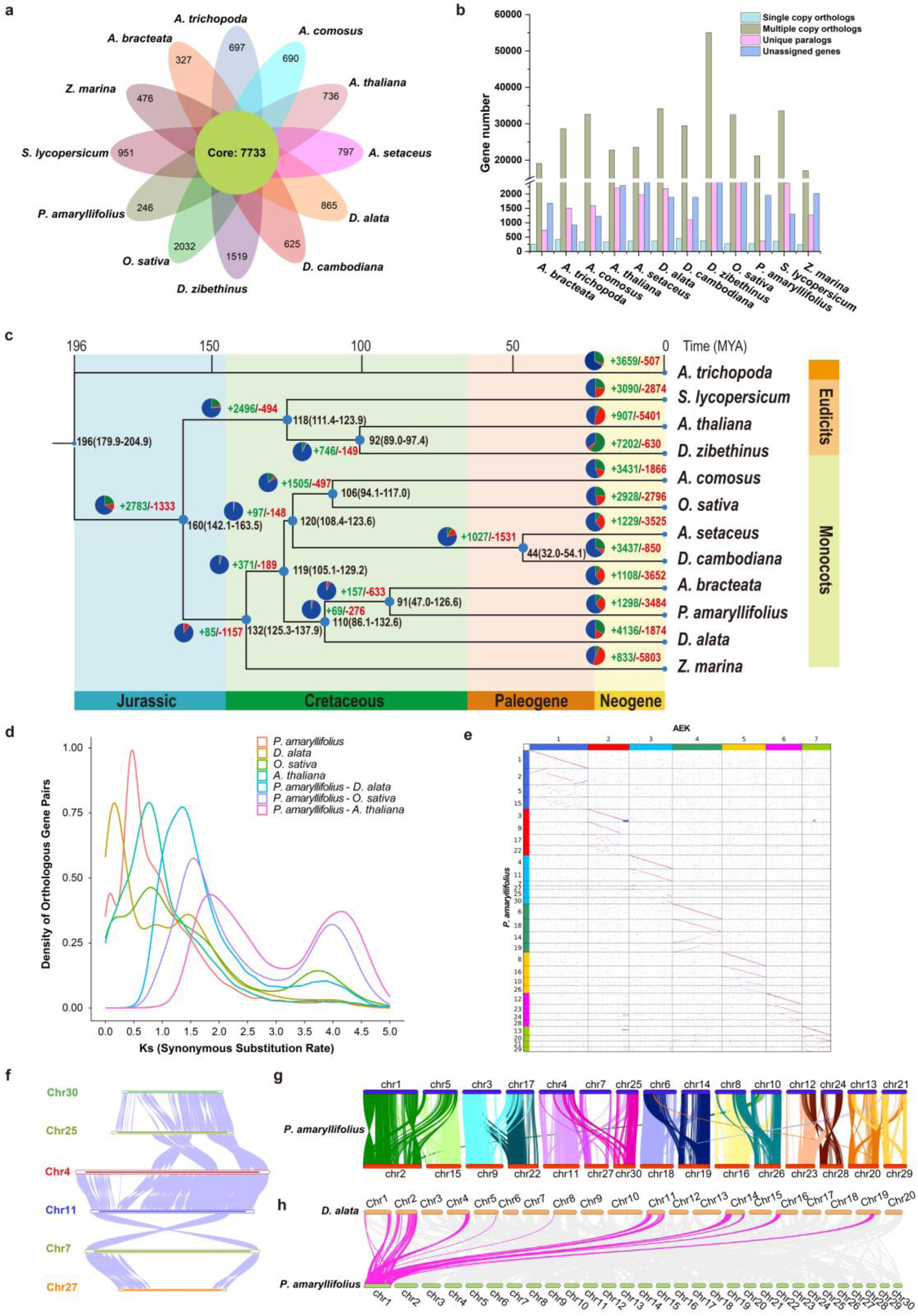
Comparative genomic analysis of *P. amaryllifolius* and 11 other species. (a) Venn diagram representing the distribution of shared gene families among *P. amaryllifolius* and 11 other species; (b) Statistics of homologous genes among species; (c) Phylogenetic tree, divergence time, expansion and contraction of gene families among species. Inferred divergence times (million years ago, Mya) are denoted in black at each node; (d) Synonymous substitutions per synonymous site (Ks) analysis of paralogs and orthologs of *P. amaryllifolius* and other representative species; (e) The karyotype evolution of *P. amaryllifolius* from ancestral eudicot karyotypes (AEK); (f) Syntenic relationships between homologous chromosomes chr4, chr7, chr11, chr25, chr27, and chr30; (g) Gene synteny within the *P. amaryllifolius* genome; (h) Syntenic patterns between *P. amaryllifolius* and *D. alata*.

To infer the evolutionary position of *P. amaryllifolius*, a phylogenetic tree was constructed via concatenated sequence alignment of 241 single-copy orthologs shared across *P. amaryllifolius* and the other 11 representative plant species. The phylogenetic tree confirmed that *P. amaryllifolius* clusters with *A. bracteatus* and *D. alata*, and the estimated divergence time between *P. amaryllifolius* and *A. bracteatus* was ∼91 Mya (Fig. 2c). Gene family contraction and expansion analysis showed that 1,284 orthologous groups (1,961 genes) were expanded in *P. amaryllifolius*, whereas 3,484 orthologous groups (3,855 genes) were contracted (Fig. 2c). KEGG enrichment analysis revealed that the expanded orthologous groups were enriched in pathways including phenylpropanoid biosynthesis, monoterpenoid biosynthesis, brassinosteroid biosynthesis, and terpenoid and polyketide metabolism, all of which are implicated in plant aroma biosynthesis (Table S18, Table S19). A total of 111 genes potentially under positive selection were identified based on Ka/Ks ratio estimates, with the majority enriched in plant-pathogen interaction, environmental adaptation, and organismal systems pathways (Table S20, Table S21).

To further investigate genome evolution and expansion, WGD events were inferred based on the Ks distribution among collinear gene pairs. Ks analysis for *P. amaryllifolius* revealed two distinct peaks at ∼0.08 and ∼0.46, indicating two independent WGD events following species divergence (Fig. 2d). These events were estimated to have occurred at ∼13 Mya and ∼74 Mya, respectively. The presence of one-versus-three syntenic blocks provided additional support for two rounds of WGD in the *P. amaryllifolius* genome (Fig. S10). The karyotype evolution of *P. amaryllifolius* from AEK revealed that most ancestral chromosomes experienced two rounds of WGD, while one ancestral chromosome underwent an additional round of WGD, and no obvious chromosomal rearrangements were detected (Fig. 2e-g). Collinearity analysis further revealed high interspecific conservation between *P. amaryllifolius* and *D. alata*, while genes located on duplicated chromosomes in *P. amaryllifolius* exhibited structural variations, including tandem duplication, translocation, and deletion (Fig. 2g, h). LTR insertion time analysis revealed that *P. amaryllifolius* exhibited moderate transposon activity during the relatively recent evolutionary period (∼0.14 Mya) and maintained a stable transposon activity pattern across long-term evolution (Fig. S11). These findings delineate a complex evolutionary history shaped by WGD, and the selective expansion of metabolism-associated gene families.

### 3.5 Differential accumulation of aroma-related metabolites under shading treatments

To examine the effects of shading on aroma biosynthesis in *P. amaryllifolius*, the characteristic aroma compound 2-AP was initially quantified in leaf samples and volatile aromas of the species’ leaves (Fig. 3a). Under 50% saturating light, the total 2-AP content in leaves was significantly higher at 1 and 3 weeks than under saturating light (100%) (*p < 0.01*), but no significant difference was observed at 7 weeks (Fig. S12). However, no significant variations in 2-AP levels were detected in the volatile aromas between the two light treatments, suggesting that differences in *P. amaryllifolius* leaf aromatic perception may predominantly stem from other aroma-active compounds. Subsequently, volatile metabolite profiling was conducted on leaf samples grown under 50% saturating light and saturating light conditions for 1, 3, and 7 weeks (designated as 50_1w, 50_3w, 50_7w, 100_1w, 100_3w, and 100_7w). The PCA revealed clear sample separation, with leaf metabolite profiles displaying more distinct separation across growth durations than light treatments (Fig. 3b). A total of 1,599 VOCs were detected and classified into 15 categories (Fig. 3c). Terpenoids (18.76%) were the predominant class, followed by esters (16.51%), ketones (11.76%), heterocyclic compounds (10.19%), and alcohols (9.26%) (Fig. 3c). As shown in Figs. 3c and 3d, significant differences in VOC profiles existed among the six groups, with both total abundance and individual compound levels varying substantially with growth durations and light treatments (Fig. 3d, e, Table S22). Notably, terpenoids, ketones, and alcohols exhibited the highest abundances (up to 54.65 μg/g, 41.11 μg/g, and 36.38 μg/g, respectively) in 100_7w leaves (Fig. 3d). Relative odor activity values (rOAVs) of the VOCs increased with leaf growth duration and were significantly higher under saturating light conditions than under 50% saturating light (Fig. 3f, Table S23). Specifically, β-ionone (terpenoid), dihydro-2-methyl-3(2H)-furanone (ketone), ethyl 3-cyclohexenecarboxylate (ester), 1,3,5-trithiane (heterocyclic compound), α,α,4-trimethyl-3-cyclohexene-1-methanethiol (alcohol), and (E,Z)-2,6-nonadienal (aldehyde) were the most odor-active. Collectively, these compounds serve as key contributors to the characteristic leaf aroma of *P. amaryllifolius*.

**Fig. 3.**
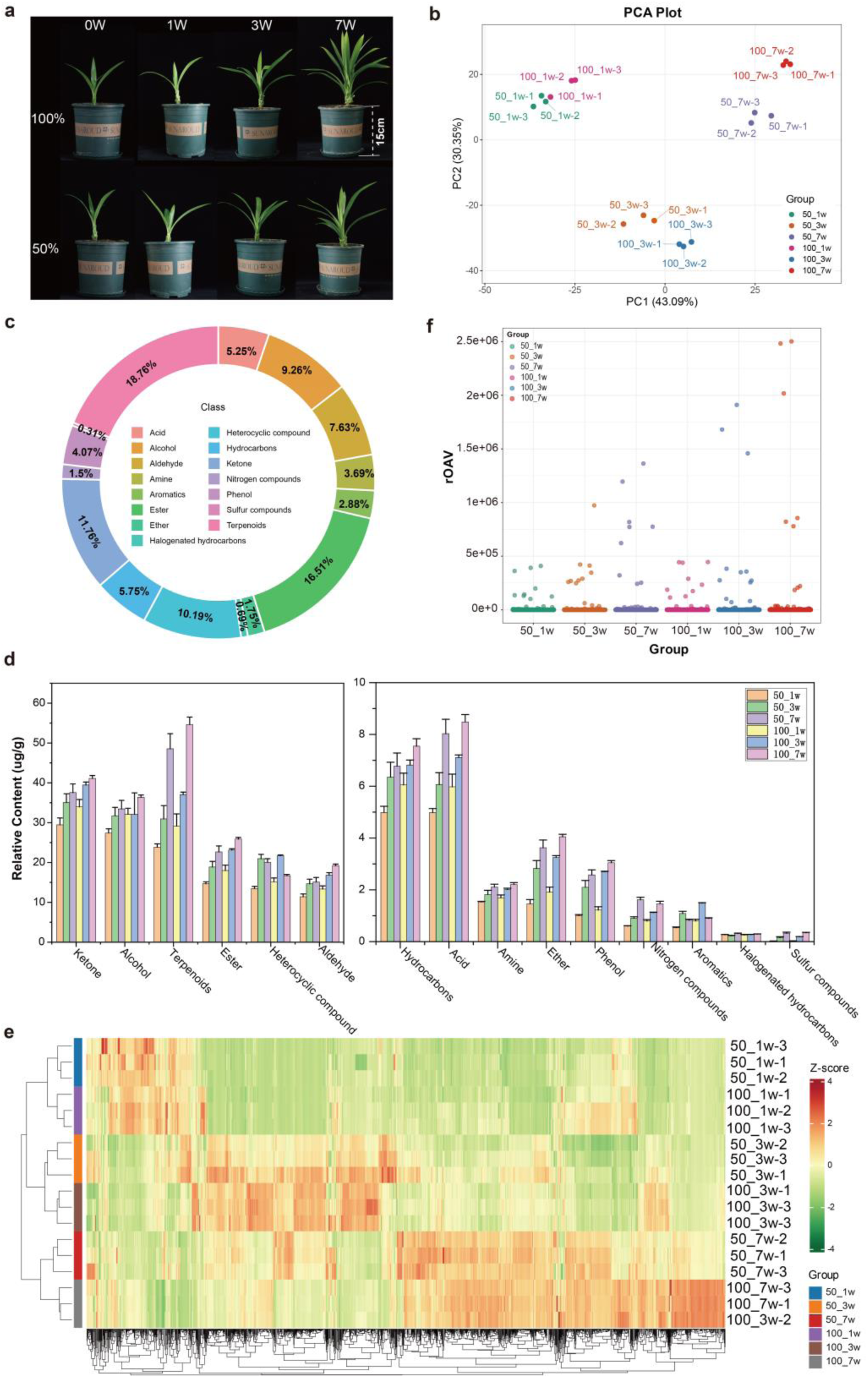
Comparison of volatile organic compound (VOC) profiles in leaf samples of *P. amaryllifolius* under 50% saturating light and saturating light at 1, 3, and 7 weeks post-treatment. (a) *P. amaryllifolius* seedlings grown under 50% saturating light and saturating light (100%) at 0, 1, 3, 7 weeks; (b) PCA of VOCs detected in leaf samples; (c) Classification and relative proportion of VOCs detected in leaf samples; (d) Total relative concentrations of major VOCs classes in the leaf samples; (e) Heatmap of relative contents of all the detected VOCs in leaf samples; (f) Scatter plot of relative odor activity values (rOAV) of aroma-active VOCs across different groups.

To identify key metabolites underlying shading-dependent aroma variation, a comparative analysis was conducted across the six groups stratified by shading conditions and growth durations to pinpoint compounds influencing aroma biosynthesis. A total of 136, 177, and 299 differentially accumulated volatiles (DAVs) were identified in the 50_1w vs. 100_1w, 50_3w vs. 100_3w, and 50_7w vs. 100_7w comparisons, respectively (Fig. 4a). Notably, the number of both up- and down-regulated DAVs increased with leaf growth progression across the three comparisons. KEGG analysis revealed that these DAVs were enriched in VOC biosynthesis pathways, particularly monoterpenoid, sesquiterpenoid and triterpenoid biosynthesis (Fig. 4b). Flavor characteristic analysis of DAVs in the three comparison groups indicated that leaf aromas were mainly characterized by sweet, fruity, green, and woody notes (Fig. 4c). A total of 142 odor-associated terpenoid DAVs were identified, most of which accumulating at higher levels under saturating light at 7 weeks (Fig. 4d). Venn diagram analysis identified 18 shared DAVs, 9 of which were functionally involved in aroma biosynthesis. Compared with those under saturating light, five shared DAVs (2-methyl-2-heptanal, 2-ethylhexanal, acetylpyrazine, (Z)-4-acetoxy-3-penten-2-one, and 2-ethylhexanol) were more abundant under 50% saturating light, contributing to green, herbal, roasted, and nutty aromatic notes. In contrast, the remaining four DAVs (paroxypropione, (E)-α-bisabolene, (Z)-γ-bisabolene, and α-bisabolene) showed lower accumulation under 50% saturating light, responsible for sweet, woody, and complementary nuances. Notably, the three bisabolene isomers are terpenoid signature metabolites defining plant specific flavors. Additionally, several off-flavor metabolites, including 4-pyridinecarboxaldehyde, (S)-N-methyl-2-pyrrolidinemethanamine, and aniline, were elevated under 50% saturating light. The differential accumulation patterns of these aroma-active DAVs and off-flavor compounds likely explain the specific flavor deterioration observed under 50% saturating light conditions.

**Fig. 4.**
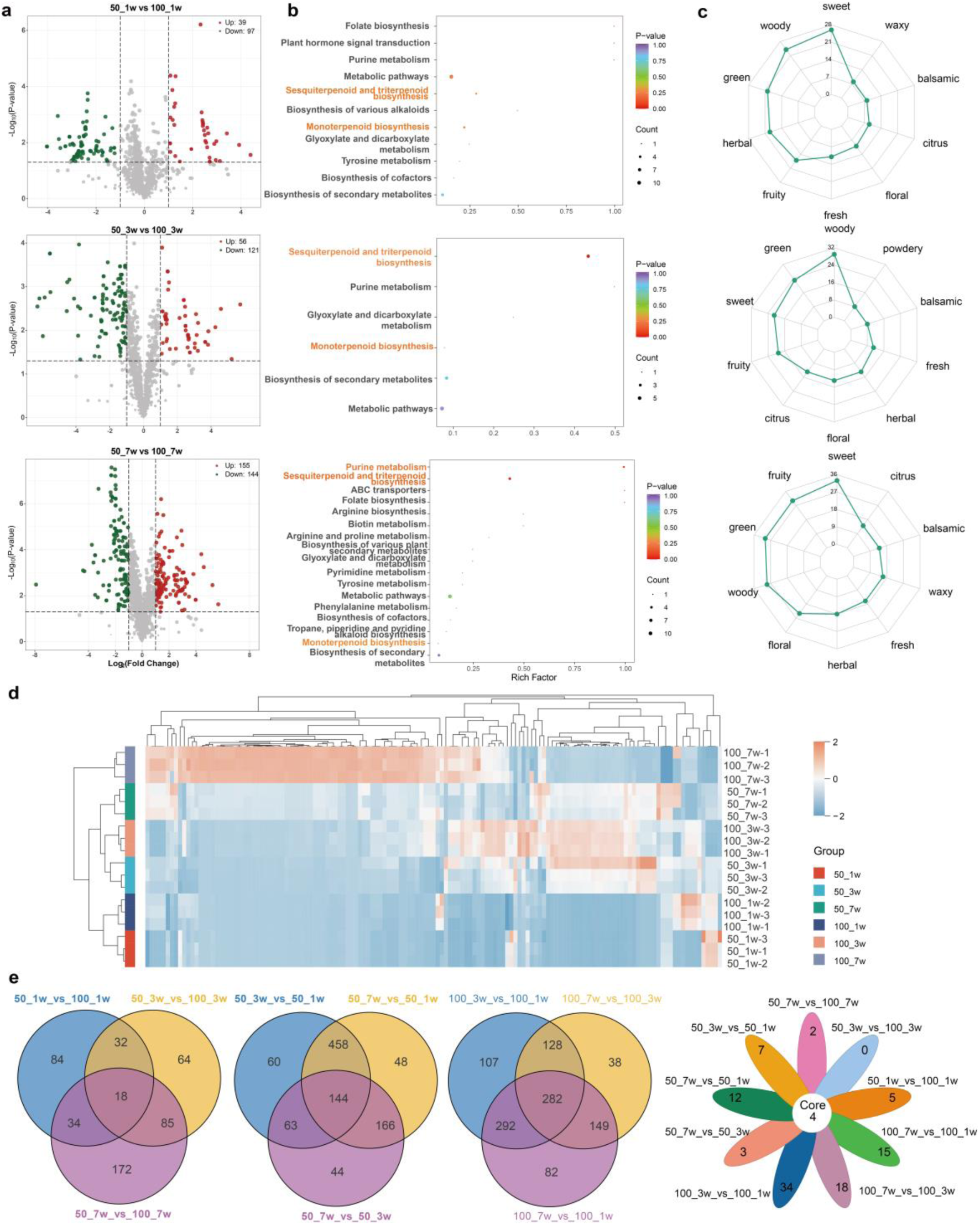
Differentially accumulated volatiles (DAVs) in leaf samples of *P. amaryllifolius* under 50% saturating light and saturating light at 1, 3, and 7 weeks post-treatment. (a) Volcano plots of DAVs in the three comparison groups; (b) KEGG analysis of DAVs in each comparison group; (c) Odor profiles of DAVs across the three comparison groups; (d) Heatmap of odor attribute-associated terpenoid DAVs across all comparison groups; (e) Venn diagram representing the overlap of shared DAVs among the three comparison groups.

Further, pairwise comparisons were performed within the 50% saturating light (50_1w, 50_3w, 50_7w) and saturating light groups (100_1w, 100_3w, 100_7w) separately, along the growth time gradient. More DAVs were detected among time-point comparisons in each group, with KEGG analysis showing that sesquiterpenoid and triterpenoid biosynthesis, phenylpropanoid biosynthesis, monoterpenoid biosynthesis, and other metabolic pathways may play crucial roles in the temporal regulation of aroma accumulation (Fig. 4e, Fig. S13 and Fig. S14). Intra-group flavor analysis indicated that sweet, green, fruity, and woody notes were primarily dominant under 50% saturating light, whereas the order was sweet, fruity, green, and woody under saturating light. This observation suggested that the inter-regime differences in leaf aroma may be associated with variations in the relative abundances of green and fruity notes. In addition, four DAVs were shared across all intra-group comparisons: three bisabolene derivatives ((E)-α-bisabolene, (Z)-γ-bisabolene, α-bisabolene) and one ketone (paroxypropione). These results collectively confirmed that terpenoids play a pivotal role in shaping the leaf aroma of *P. amaryllifolius*.

### 3.6 Transcriptomic combined with WGCNA analysis characterizes genes underpinning terpenoid-mediated aroma biosynthesis

To investigate the biosynthesis of VOCs and the molecular mechanisms underlying leaf aroma formation in *P. amaryllifolius*, RNA sequencing was conducted on leaf samples collected at 1, 3, and 7 weeks under 50% saturating light and saturating light conditions. Across the three developmental stages, numerous DEGs were detected between the two light regimes (Figs. 5a–c). GO and KEGG enrichment analyses revealed that these DEGs were significantly enriched in light signaling, hormone response, terpene biosynthesis, and secondary metabolic pathways linked to the above processes (Fig. 5d, Fig. S15 and Fig.S16). Among all 4,221 identified DEGs, 132 genes were shared across the three comparison groups (50_1w vs. 100_1w, 50_3w vs. 100_3w, and 50_7w vs. 100_7w). These shared DEGs were also significantly enriched in pathways related to environmental information processing, genetic information processing, light signal, and aroma biosynthesis (Fig. 5e and Fig.S17). Notably, several key TFs involved in aroma biosynthesis (e.g., MYB, bHLH) and key terpene biosynthesis enzymes (e.g., TPS (*Pam1100361*, *PaTPS18*), carotenoid cleavage dioxygenase 4 (CCD4, *novel.33*)) were identified. The consistent identification of TPS and CCD4 between 50% saturating light and saturating light comparisons further underscored their central roles in regulating leaf aroma accumulation in *P. amaryllifolius*.

**Fig. 5.**
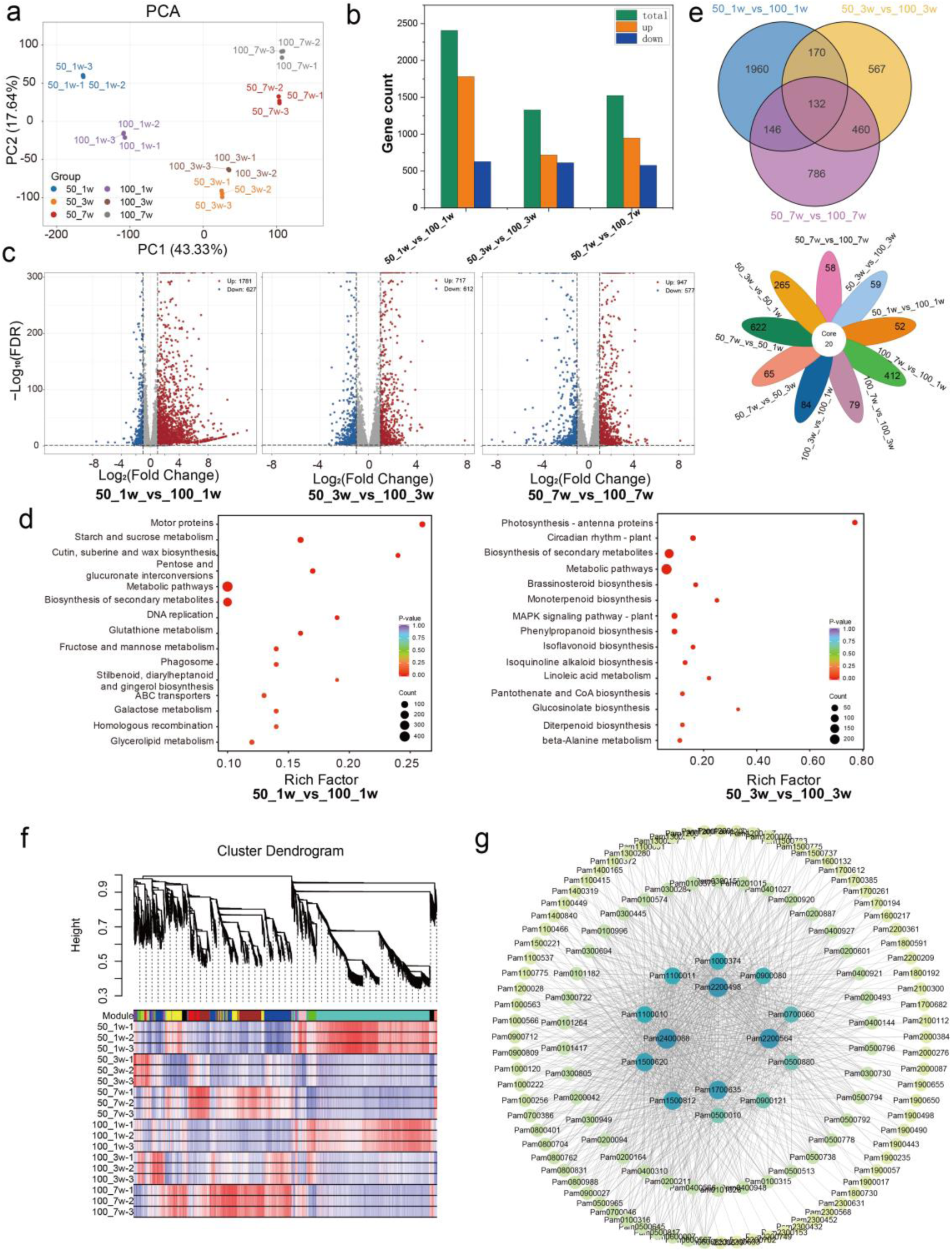
Transcriptomic and WGCNA analysis of *P. amaryllifolius* leaf samples under 50% saturating light and saturating light at 1, 3, and 7 weeks post-treatment. (a) PCA of transcriptome data from leaf samples; (b) Number of differentially expressed genes (DEGs) in the comparison groups; (c) Volcano plots of DEGs in the three comparison groups; (d) Top 15 enriched GO terms and KEGG pathways of DEGs across the comparison groups; (e) Venn diagram representing the overlap of shared DEGs; (f) Hierarchical clustering tree of co-expression modules based on WGCNA; (g) Co-expression network of hub-genes in the “yellow” module.

To further uncover the regulatory mechanisms underlying leaf aroma biosynthesis, WGCNA was performed and identified fifteen distinct gene co-expression modules via hierarchical clustering dendrograms (Fig. 5f). Notably, fifteen classes of VOCs, including significantly accumulated terpenoid DAVs, exhibited strong positive correlations with distinct modules, respectively (Fig. S18). The “brown” module showed a robust correlation with terpenoid VOC accumulation, and *Pam0200950* was identified as a hub gene of this module, encoding core rate-limiting enzyme (E)-4-hydroxy-3-methylbut-2-enyl-diphosphate synthase (*HDS*) of MEP pathway (Fig. S18 and Fig. S19). The three shared DAV terpenoids ((E)-α-bisabolene, (Z)-γ-bisabolene, and α-bisabolene) were significantly correlated with the “yellow” module, which harbored 565 DEGs (Fig. S18). The hub genes of this module were *Pam2400088* (*PaTPS19*), *Pam1700635* (encoding solanesyl-diphosphate synthase), *Pam2200498* (gibberellin 2β-dioxygenase), and *Pam2200564* (all-trans-nonaprenyl-diphosphate synthase), which directly or indirectly participated in terpenoid biosynthesis, carotenoid biosynthesis, and terpenoid backbone biosynthesis (Fig. 5g and Fig. S20). Collectively, these results highlight the central role of coordinated terpenoid pathway regulation in shaping the shading-responsive aroma phenotype of *P. amaryllifolius*.

### 3.7 Identification of structural genes in the terpenoid biosynthesis pathway

Given that monoterpenoids and sesquiterpenoids are the primary terpenoid volatile components in *P. amaryllifolius*, the structural genes involved in terpenoid biosynthesis were systematically identified. The core terpenoid precursors, isopentenyl diphosphate (IPP) and dimethylallyl diphosphate (DMAPP), are synthesized via the cytosolic MVA pathway and plastidial MEP pathway, respectively. While most terpenoid biosynthesis genes were conserved in *P. amaryllifolius*, the HMGR genes (encoding 3-hydroxy-3-methylglutaryl-CoA reductase) and DXS genes (encoding 1-deoxy-D-xylulose 5-phosphate synthase) have undergone significant expansion compared to their orthologs in *A. thaliana*, *O. sativa*, and its phylogenetically close relative *D. alata* (Table S24). Under 50% saturating light and saturating light conditions, structural genes associated with terpenoid biosynthesis in both the MVA and MEP pathways exhibited differential expression (Fig. 6a). In the MVA pathway, most DEGs were significantly upregulated after 1 week of 50% saturating light treatment compared to that under saturating light. In contrast, DEGs in the MEP pathway displayed distinct temporal expression patterns, with their expression levels gradually up-regulated from 1 to 3 weeks and further to 7 weeks of treatment.

**Fig. 6.**
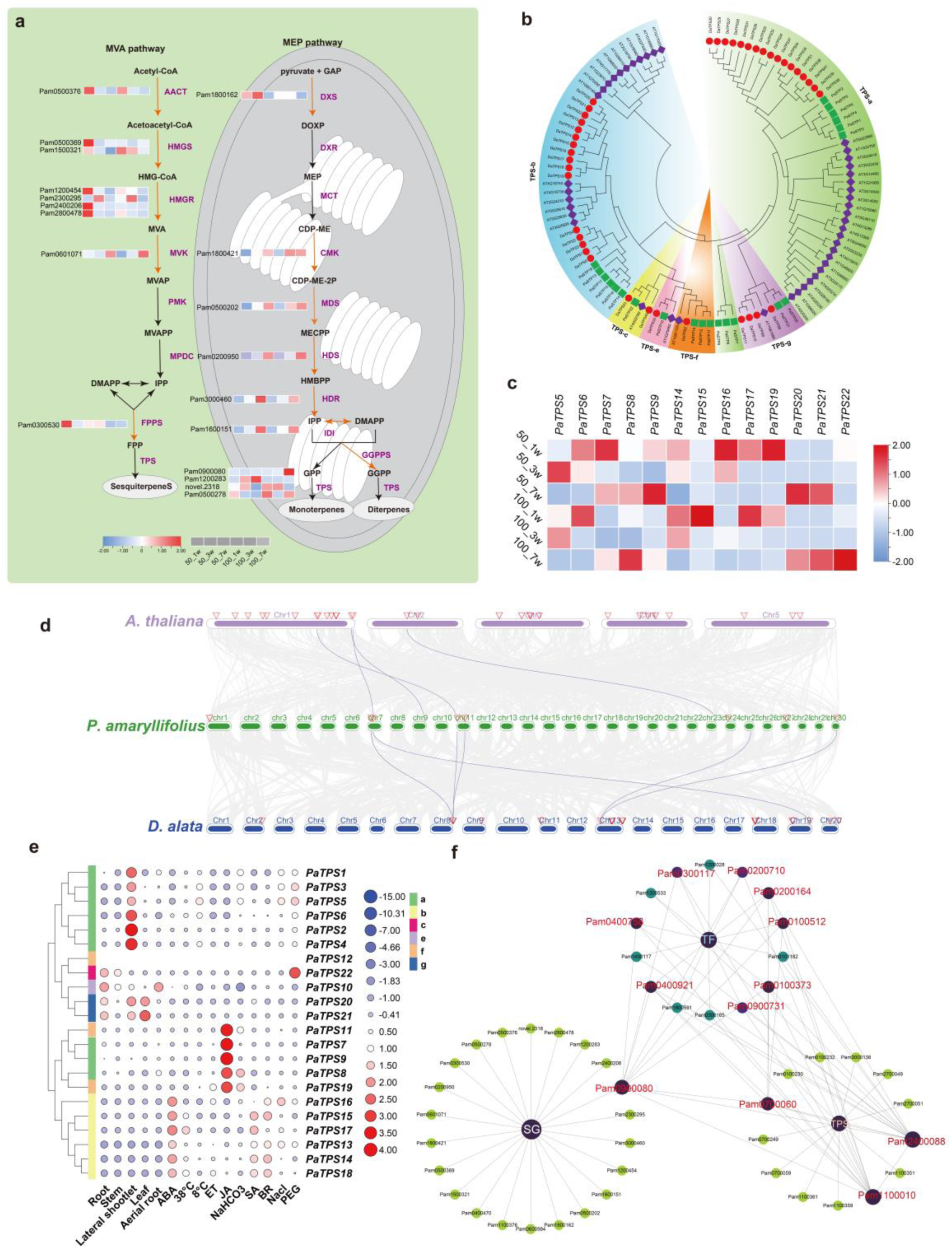
Identification of terpenoid biosynthesis genes and terpene synthase (TPS) family members in *P. amaryllifolius*. (a) Identification of structural genes in mevalonate (MVA) and methylerythritol 4-phosphate (MEP) pathways (terpenoid biosynthesis) in *P. amaryllifolius*; (b) Phylogenetic tree of TPS family members from *P. amaryllifolius*, *A. thaliana* and *D. alata*; (c) Expression heatmap of the TPS gene family in *P. amaryllifolius* leaf samples under 50% saturating light and saturating light at 1, 3, and 7 weeks post-treatment; (d) Collinearity analyses of TPS family members among *P. amaryllifolius*, *A. thaliana* and *D. alata*; (e) Expression heatmap of the *P. amaryllifolius* TPS gene family across different tissues and treatments; (f) Co-expression network of terpenoid biosynthesis structure genes (SG), TPS genes and transcription factor DEGs.

TPS catalyze the final step of terpenoid biosynthesis, converting farnesyl diphosphate (FPP), geranyl diphosphate (GPP), and geranylgeranyl diphosphate (GGPP) into diverse terpenoids. In this study, 22 TPS family members were systematically identified from the *P. amaryllifolius* T2T genome and were clustered into TPS-a, TPS-b, TPS-c, TPS-e, TPS-f, and TPS-g subfamilies (Fig. 6b). Among these, 9 TPS-a and 6 TPS-b members dominated the family and were primarily responsible for sesquiterpenoid and monoterpenoid biosynthesis, respectively (Table S25). A total of 5 TPS-a and 4 TPS-b genes showed significant expression differences between 50% saturating light and saturating light (Fig. 6c). Additionally, genes belonging to TPS-c, TPS-g, and TPS-f subfamilies were differentially expressed under the two light conditions, and these expression variations collectively contributed to terpenoid diversity and aroma profile formation. Notably, *P. amaryllifolius* harbored significantly fewer TPS genes than *A. thaliana* and its phylogenetically close relative *D. alata*, indicating notable TPS family contraction. Synteny analysis revealed substantial conservation, losses and functional divergence of TPS genes (Fig. 6d). Transcriptomic data showed that all PaTPS genes, except *PaTPS12,* exhibited differential expression across tissues, phytohormone treatments, and stress conditions, with a subset of TPS genes tightly regulated by ABA, JA, SA, and BR (Fig. 6e).

Co-expression network analysis demonstrated robust co-expression among terpenoid biosynthesis-related structural genes, TPS genes, and diverse TFs. Eight TFs (*Pam0100373*, *Pam0100512*, *Pam0200164*, *Pam0200710*, *Pam0300117*, *Pam0400758*, *Pam0400921*, *Pam0900731*) showed strong positive co-expression with three TPS genes (*PamTPS9*, *PamTPS12*, *PamTPS19*) and one structural gene (*Pam0900080*) (*r² > 0.8* and *p < 0.001*) (Fig. 6f). The TF binding sites were also identified in the promoters of these target genes (Fig. S21, Table S26). Notably, gene-metabolite correlation analysis revealed that key differential terpenoids, including (E)-α-bisabolene, (Z)-γ-bisabolene and α-bisabolene, exhibited strong positive correlations with the aforementioned TFs, structural genes and TPS genes (*r² > 0.8* and *p < 0.001*) (Fig. S22). In addition, *Pam0900731*, encoding a Heat Shock Factor (HSF), exhibited consistently strong positive co-expression with structural genes and TPS genes in both co-expression network analysis and gene-metabolite correlation analysis, suggesting it may act as the core TF regulating the synthesis of terpenoid DAVs. Furthermore, the expression patterns of terpenoid biosynthesis-related structural genes, TPS genes, and TFs were experimentally validated via qRT-PCR, with the majority of these genes showing expression profiles consistent with transcriptomic data (Fig. 7). Finally, based on these integrated findings, we proposed a regulatory model illustrating how shading modulates leaf aroma biosynthesis in *P. amaryllifolius* via a coordinated network mediated by the MVA/MEP pathways, TPS family members, and key TFs (Fig. 8).

**Fig. 7.**
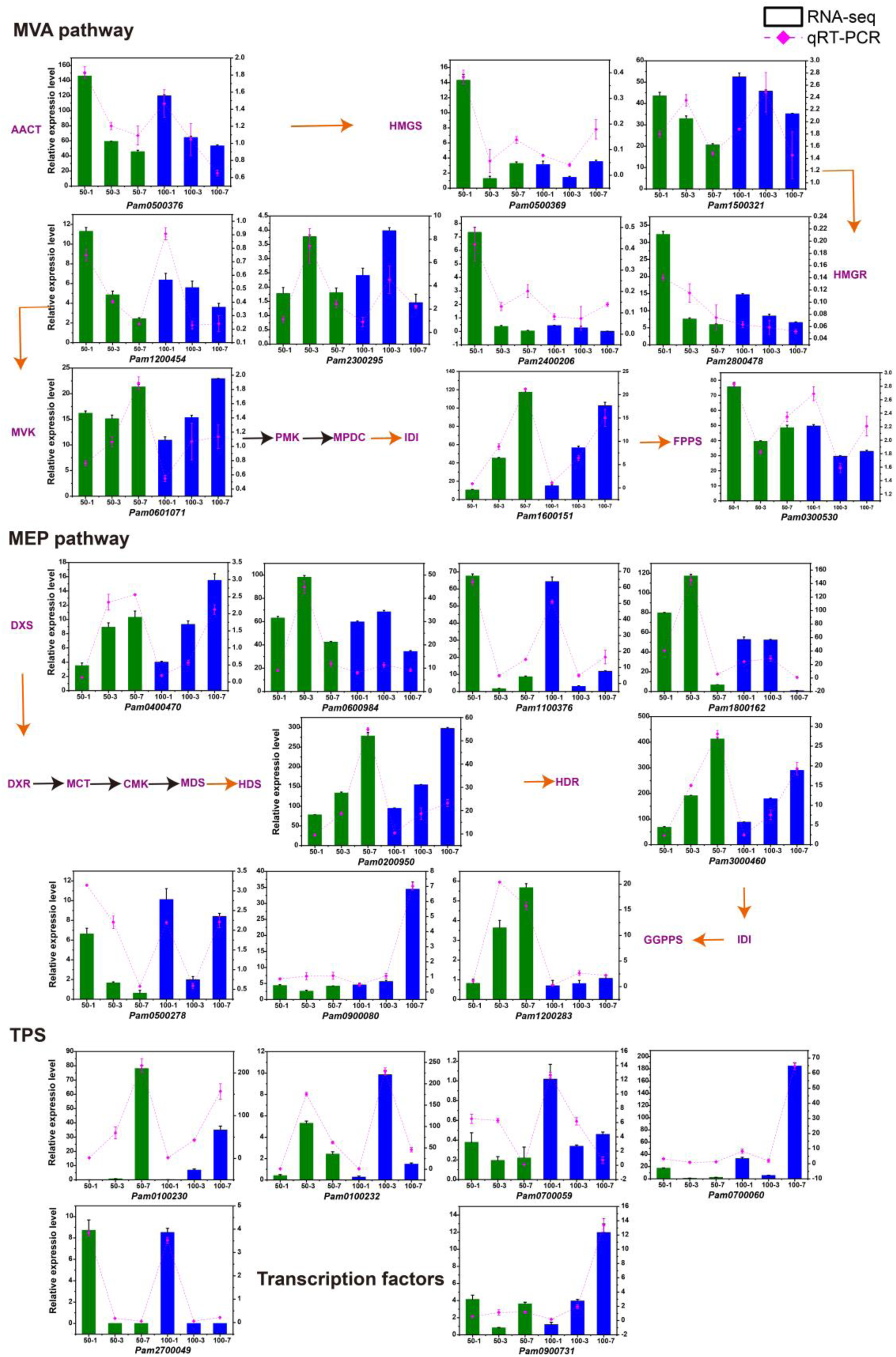
Validation of expression levels of structural genes in the mevalonate (MVA) and methylerythritol 4-phosphate (MEP) Pathways, TPS genes, and transcription factors in leaf samples of *P. amaryllifolius* under 50% saturating light and saturating light at 1, 3, and 7 weeks post-treatment. Bar charts indicate FPKM-based expression levels derived from RNA-seq data, while line plots represent the relative expression levels quantified via qRT-PCR.

**Fig. 8.**
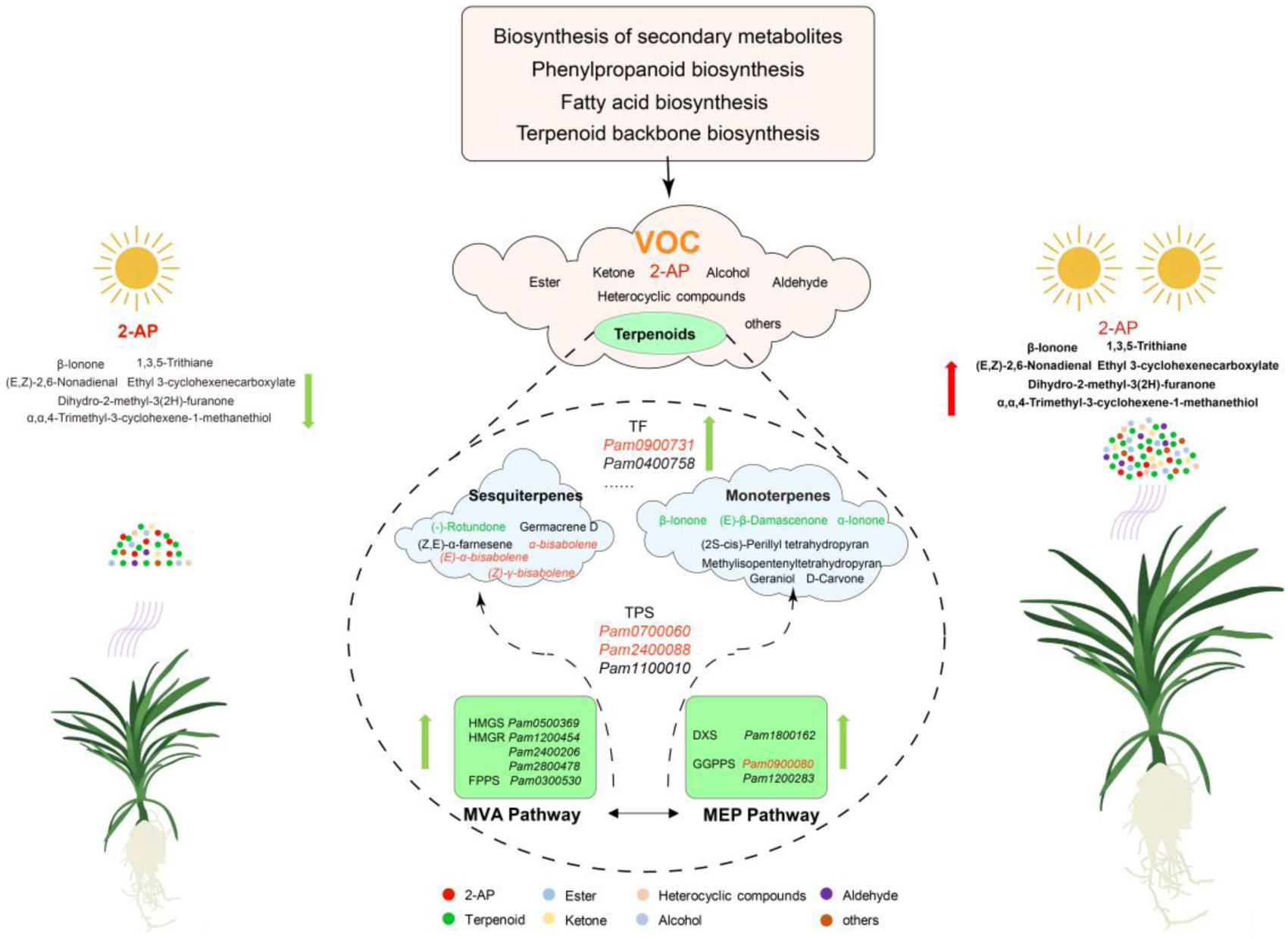
Schematic model illustrating the regulation of differentially accumulated volatiles (DAVs) in *P. amaryllifolius* leaves under 50% saturating light and saturating light conditions. Numerous DAVs, including terpenoids, esters, ketones, heterocyclic compounds, and aldehydes, were detected in *P. amaryllifolius* leaves, with terpenoids identified as the primary contributors to leaf aroma. Key mevalonate (MVA) and methylerythritol 4-phosphate (MEP) pathway structural genes, TPS family members, and terpenoid-related transcription factors are illustrated, forming a coordinated network regulating shading-responsive aroma variation in *P. amaryllifolius*.

## Discussion

A complete, high-quality, and accurately annotated reference genome serves as a foundational resource for functional genomics, molecular breeding, evolutionary studies, and genomic selection in crops (Varshney et al., 2014; Yadav et al., 2025). *P. amaryllifolius* is a multifaceted valuable plant, renowned for its unique fragrance, nutritional benefits, and medicinal applications. However, the only previously reported genome assembly for this species was highly fragmented and lacked chromosome-level assembly contiguity, severely limiting genomic and functional investigations (Sidek et al., 2025). Here, we presented the first T2T gap-free genome assembly of *P. amaryllifolius* cultivar ‘Taishan Banlan’, which consisted of 30 chromosomes with a contig N50 of 17.00 Mb, marking a substantial advancement over the prior assembly (1,448 contigs, contig N50 of 9.02 Mb). Complete resolution of centromeres and telomeres, coupled with rigorous quality validation and reliable genome annotation, collectively confirmed that this assembly achieves high-quality standards and provided a key milestone for subsequent genomic investigations of *P. amaryllifolius*.

*P. amaryllifolius* is a shade-preferring aromatic plant with a low light saturation point, cultivated in the understories of tropical economic forests (Yan et al., 2025; Zhang et al., 2023). Previous studies showed that when the shading degree ranged from 30% to 60%, *P. amaryllifolius* maintained favorable growth performance and a high content of 2-AP (Tang et al., 2020). In this study, we confirmed that shading facilitated leaf 2-AP at the early stage (1 week post-treatment) but did not elicit significant changes in volatile 2-AP levels, which was likely due to the pre-existing high basal concentration of 2-AP in leaves. Beyond its role in regulating 2-AP, we identified that shading played a key role in suppressing the synthesis of volatile compounds, especially terpenoids, and consequently altered the leaf aroma profile. Terpenoids are core aromatic contributors in many fragrant plants, including water dropwort (Feng et al., 2025), strawberries (Fang et al., 2024), and lotus (Chen et al., 2025). Comparative metabolomic analysis of *P. amaryllifolius* leaves under 50% saturating light and saturating light identified three bisabolene isomers ((E)-α-bisabolene, (Z)-γ-bisabolene, and α-bisabolene) as the primary shared DAVs. Bisabolene is characterized by a warm, sweet-spicy-balsamic odour and is known to enhance aroma stability and persistence (Li et al., 2023). Collectively, the reduced accumulation of bisabolene isomers, alongside other terpenoids, esters, ketones, and other related compounds, under 50% saturating light likely underpins the distinct differences in leaf aroma observed between the 50% saturating light and saturating light conditions.

In plants, monoterpenes and sesquiterpenes are synthesized via the plastidial MEP and the cytosolic MVA pathways (Ramya et al., 2020). Leaf transcriptomic and WGCNA analyses identified *Pam0200950* as a hub gene in terpenoid biosynthesis, encoding the MEP pathway HDS, and *Pam2400088* as a bisabolene-specific hub gene encoding a member of TPS subfamily f. Furthermore, compared with saturating light conditions, genes up-regulated under 50% saturating light were primarily enriched in the MVA pathway at 1 week post-treatment. The TPS gene family, which serves as a key component in the final step of terpenoid biosynthesis, determines the abundance and diversity of terpenoids across plant species (Bao et al., 2020; Zheng et al., 2024). In this study, 22 TPS genes were identified in *P. amaryllifolius*, and 21 of them were expressed and exhibited hormone-responsive expression patterns. Relative to *A. thaliana* and *D. alata*, the TPS gene family of *P. amaryllifolius* displayed a notable contraction, an evolutionary feature shared with Lotus (Chen et al., 2025). Conversely, several other fragrant plant species, including *O. javanica* (Feng et al., 2025), have undergone TPS family expansion, suggesting that *P. amaryllifolius* may have evolved a specialized terpenoid profile. WGCNA and gene-metabolite correlation network analyses identified co-expression relationships among TF genes, structural genes, and TPS genes, indicating a complex regulatory network for terpenoid biosynthesis. Interestingly, a previous study showed that same subfamily TPS proteins, have different subcellular localizations and functions in terpenoid biosynthesis (Chen et al., 2025; Li et al., 2024). Thus, future in-depth studies are required to clarify the terpenoid biosynthesis pathway and its role in the leaf aroma of *P. amaryllifolius*.

Transcriptional regulation is critical for terpenoid biosynthesis, and multiple TF families (e.g., AP2, NAC, bZIP, WRKY, HSF, MYB, bHLH) have been widely reported to participate in this process (Jin et al., 2025). Consistent with this, an HSF transcription factor (Pam0900731) was identified to be strongly associated with three key terpenoid DAVs, three TPS genes (*PamTPS9*, *PamTPS12*, *PamTPS19*) and one structural gene (*Pam0900080*). This finding implies that HSF may act as a core terpenoid regulator by modulating TPS genes or structural genes. Notably, accumulating evidence has uncovered diverse regulatory modes of HSF in terpenoid metabolism, providing valuable comparative frameworks for deciphering HSF-mediated regulation in *P. amaryllifolius*. For instance, in *Andrographis paniculata*, ApHSFB2b forms a protein complex with ApMYC2, which regulates the TPS gene *ApCPS1* via JA signaling to promote andrographolide accumulation (Huang et al., 2025). Intriguingly, our study found that *PamTPS9* was significantly up-regulated by exogenous JA, a pattern analogous to *ApCPS1* regulation in *A. paniculata*, hinting at potential crosstalk between HSF, JA signaling, and TPS genes in *P. amaryllifolius* (Huang et al., 2025). In contrast, *PtHSF1* in *Phaeodactylum tricornutum* directly regulates the key MEP pathway gene *DXS* to mediate fucoxanthin biosynthesis (Song et al., 2023), representing a distinct direct regulatory mode of HSF in isoprenoid metabolism. Meanwhile, promoter sequence analysis of *PamTPS9* revealed abundant TF binding sites, including heat shock elements (HSEs), MYC, and MYB sites, further supporting the potential for combinatorial TF regulation of this key TPS gene. Despite these insightful clues, the specific regulatory mechanism by which HSF modulates terpenoid biosynthesis in *P. amaryllifolius* remains elusive and merits detailed mechanistic investigation in future studies.

## Data availability

The raw whole-genome sequencing data of *P. amaryllifolius* are available at the National Genomics Data Center (NGDC), China National Center for Bioinformation (CNCB; https://ngdc.cncb.ac.cn/), and can be accessed via the project accession number PRJCA044941.

## CRediT authorship contribution statement

Zhijun Xu: Conceptualization, Data curation, Formal analysis, Resources, Visualization, Writing-original draft, Writing-review & editing. Xuejiao Zhang: Funding acquisition, Investigation, Methodology. Yuzhan Li: Validation, Visualization, Writing-review & editing. Sheng Zhao: Validation, Visualization, Writing-review & editing. Qibiao Li: Investigation, Methodology. Jiannan Zhou: Data curation, Formal analysis, Methodology, Software. Huan Ouyang: Conceptualization, Resources, Software, Validation. Xiaowen Hu: Conceptualization, Data curation, Formal analysis, Software, Visualization, Writing-review & editing.

## Acknowledgements

This work was supported by Central Public-interest Scientific Institution Basal Research Fund (NO. 1630102024009 and NO. 1630012025806)

